# Inhibition of lysyl oxidase-like 2 ameliorates folic acid-induced renal tubulointerstitial fibrosis

**DOI:** 10.1101/2020.12.01.406009

**Authors:** Sung-Eun Choi, Nara Jeon, Hoon Young Choi, Hyeon Joo Jeong, Beom Jin Lim

## Abstract

Tubulointerstitial fibrosis is characterized by accumulation of the extracellular matrix in the interstitium. Lysyl oxidase-like 2 (LOXL2), a member of the lysyl oxidase family, is known for promoting cancer metastasis, invasion, and stromal fibrosis in various organs. Our previous study demonstrated expression of LOXL2 in kidney podocytes and tubular epithelial cells, and the association between elevated LOXL2 and tubulointerstitial fibrosis. The present study evaluated the effect of LOXL2 inhibition using an inhibitory monoclonal antibody (AB0023) on tubulointerstitial fibrosis in a folic acid-induced tubulointerstitial fibrosis mouse model. We also evaluated the association of LOXL2 with epithelial-mesenchymal transformation related molecules *in vitro* using HK-2 cells. Our data demonstrate that AB0023 prevented the progression of tubulointerstitial fibrosis significantly, as determined by trichrome and picro-sirius red staining, as well as the total collagen assay. The mean expression of phosphorylated Smad2 and Smad4 was lower in the AB0023-treated group although it was not statistically significant. Following transforming growth factor-β (TGF-β) challenge, LOXL2-deficient HK-2 cells exhibited significantly lower expression of the mesenchymal markers vimentin and fibronectin than control HK-2 cells. In conclusion, LOXL2 inhibition ameliorates renal fibrosis through the TGF-β/Smad signalling pathway.

## Introduction

As tubulointerstitial fibrosis is a common endpoint in renal disease with no effective treatment other than dialysis, the need to understand the molecules and mechanisms involved is increasingly urgent. Histologically, tubulointerstitial fibrosis is an accumulation of the extracellular matrix (ECM) in the interstitium. ECM-producing cells are primarily activated fibroblasts [1]. Various cells such as pericytes, endothelial cells, residual fibroblasts, and tubular epithelial cells are known to be the origin of fibroblasts [2].

The epithelial-mesenchymal transformation (EMT) has been studied in cancer and benign fibrotic diseases [3]. Once acute injury is imposed on the kidney, various chemokines and growth factors cause inflammation, which in turn leads to the secretion of transforming growth factor-β (TGF-β) via release of active TGF-β from latent TGF-β-binding protein via protease cleavage [4]. TGF-β is the primary molecule responsible for EMT [5, 6], and the canonical and non-canonical pathways are the downstream pathways of TGF-β [4, 7]. The hallmark of EMT is loss of epithelial phenotypes and acquisition of mesenchymal phenotypes with activation of profibrotic genes to produce the ECM, including fibronectin and types I and III collagen [3, 5, 8].

Lysyl oxidase-like 2 (LOXL2) is a member of the lysyl oxidase family, originally known as a copper-dependent amine oxidase, that is involved in cross-linking collagen and elastin of the ECM [9]. Studies have also demonstrated additional functions for LOXL2 independent of its catalytic activity, such as organ development [10], tumour invasion [11], and EMT [12, 13].

In our previous study, we found that LOXL2 is expressed in kidney podocytes and tubular epithelial cells, and its expression is increased in the folic acid-induced murine fibrosis model [14]. In this study, we evaluated the effect and therapeutic role of LOXL2 inhibitor AB0023 on the progression of tubulointerstitial fibrosis in mice. In order to evaluate the contribution of LOXL2 in EMT, an *in vitro* study using immortalized human proximal tubular epithelial cells (HK-2 cells) was also performed.

## Methods

### Animal model of tubulointerstitial fibrosis and LOXL2 inhibition

Male CD1 mice at 8 weeks of age (Orient Bio, Inc., Seongnam, South Korea) were used in this study. The animals were housed in a facility maintained at 20°C and 12-h alternating light/dark cycles with free access to rodent chow and water. Tubulointerstitial fibrosis was induced by intraperitoneal injection of folic acid (240 μg/g body weight) [15, 16]. The folic acid solution was prepared by dissolving folic acid powder (Sigma-Aldrich) in 0.3 M NaHCO_3_. Control CD1 mice were injected intraperitoneally with the same volume of vehicle (NaHCO_3_). Urinary excretion of neutrophil gelatinase-associated lipocalin (NGAL) was measured immediately before injection and at 3 days after injection using a Mouse Lipocalin-2/NGAL Quantikine enzyme-linked immunosorbent assay (ELISA) Kit (R&D Systems) to ensure successful injection of folic acid, as manifested by a log scale increase in NGAL. The concentration of urinary NGAL was normalized by the concentration of urinary creatinine as measured by the QuantiChrom Creatinine Assay Kit (BioAssay Systems, Hayward, CA, USA). Mice without an increase in NGAL at 3 days post-folic acid injection, indicating that folic acid was not successfully injected, were omitted from the study. Ultimately, 16 mice injected with folic acid and six mice injected with vehicle were examined in this study.

To inhibit LOXL2, AB0023 (Gilead Sciences, Foster City, CA, USA), an inhibitory monoclonal antibody against LOXL2, was used. Nine of the 16 folic acid-injected mice were treated with a dosage of 15 mg/kg body weight of AB0023 at 1 week before folic acid injection and twice a week for 4 weeks afterwards. The remaining seven mice were injected with immunoglobulin G (IgG) (GS-645864, Gilead Sciences) at the same dosage and on the same time schedule as the AB0023 treatment group. Mice were sacrificed via cervical dislocation 4 weeks after folic acid or vehicle injection, and the right kidneys were harvested (Fig 1). Fresh frozen tissues were stored at −70°C for western blot analysis and collagen measurement. Additional kidney tissues were fixed in 4% formaldehyde for 24 hours at room temperature and embedded in paraffin overnight at 55-65°C using an automatic tissue processer (EFTP-FAST 360; Intelsint, Turin, Italy).

**Fig 1.**
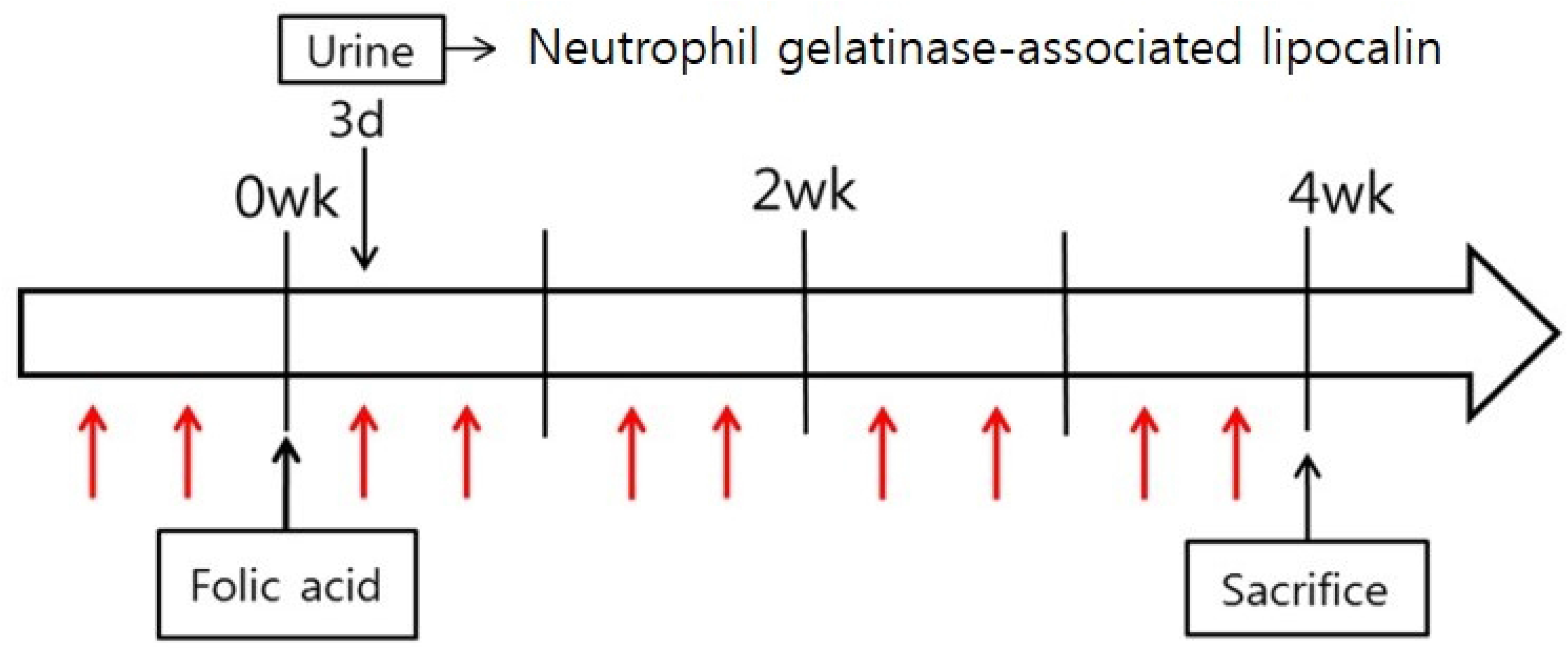
Injection protocol of folic acid, AB0023 (a monoclonal antibody against LOXL2), and control IgG in the CD1 mouse model. The red arrow indicates AB0023 or control IgG intraperitoneal injection (15 mg/kg). AB0023 or control IgG was injected 4 and 1 day before folic acid injection, and twice weekly until 4 weeks after folic acid injection. Urinary neutrophil gelatinase-associated lipocalin (NGAL) was measured 3 days after folic acid injection to ensure successful induction of renal fibrosis.

The present study was approved by the Institutional Animal Care and Use Committee of the Yonsei University Health System (Seoul, South Korea; approval number 2015-0247). All experiments involving animals were carried out in accordance with the standards set forth by the Institutional Animal Care and Use Committee of Yonsei University Health System.

### Evaluation of tubulointerstitial fibrosis

#### Semiquantitative analysis via histologic examination

Paraffin-embedded samples of the AB0023-treated, IgG-injected, and vehicle-injected groups were cut into 4-μm sections. After deparaffinization and rehydration, sections were stained with Masson trichrome and picro-sirius red. For picro-sirius red staining, sections were treated with Weigert’s Iron Hematoxylin for 8 min to stain nuclei and Direct Red 80 (Sigma-Aldrich; Merck KGaA, Darmstadt, Germany) for 1 hour at room temperature to visualize collagen before washing with 0.5% glacial acid. Slides were examined by light microscopy with or without polarisation for picro-sirius red or trichrome staining, respectively. Photos were taken serially along the cortex at 200× magnification and the area of interstitial fibrosis was measured using Image J software (version 1.50i; National Institutes of Health, Bethesda, MD, USA).

#### Quantitative analysis via total collagen analysis

The content of collagen in fresh frozen cortex was evaluated by measuring hydroxyproline using the Total Collagen Assay Kit (QuickZyme Biosciences, Leiden, Netherlands) according to the manufacturer’s guide. Briefly, samples were hydrolysed at 95°C in 6M HCl for 20 hours and then centrifuged at 13,000 rpm for 10 min. The supernatant was collected and assayed by ELISA according to the manufacturer’s protocol. Total protein in the hydrolysed sample was also measured using the Total Protein Assay Kit (QuickZyme) and the relative amount of collagen per protein was analysed.

### Renal cell culture and transfection

HK-2 cells were purchased from the American Type Culture Collection (Manassas, VA, USA) and cultured in Dulbecco’s modified Eagle’s medium (DMEM)/Nutrient Mixture F-12 (Gibco; Thermo Fisher Scientific, Waltham, MA, USA) supplemented with 10% foetal bovine serum (FBS; Gibco; Thermo Fisher Scientific). To silence LOXL2 expression at the cellular level, LOXL2 shRNA lentiviral particles (cat. no. sc-45222-v; Santa Cruz Biotechnology, Inc., Dallas, TX, USA) were transduced into HK-2 cells cultured on collagen I (2 mg/ml, cat. no. 354236; Corning Inc., Corning, NY, USA)-coated dishes[17]. HK-2 cells were treated with media containing 5 μg/ml of polybrene (cat. no. sc-134220; Santa Cruz Biotechnology, Inc.), and then LOXL2 shRNA lentiviral particles and control shRNA lentiviral particles of 1 and 2 multiplicity of infection (MOI) were added. Transfected cells were selected by selection media containing 2 μg/ml puromycin dihydrochloride (cat. no. sc-108071; Santa Cruz Biotechnology, Inc.). After confirming the decrease in LOXL2 expression by reverse transcription-quantitative polymerase chain reaction (RT-qPCR) and western blot analysis (described below), cells of 2 MOI were treated with serum-free media for 24 hours and then media containing TGF-β (20 ng/ml, R&D Systems, Minneapolis, MN, USA) for 72 hours. Other LOXL2*-*deficient cells (2 MOI) were treated with serum-free media for 24 hours and then media containing vehicle (0.1% 4mM HCl/BSA). Control shRNA lentiviral particles (cat. no. sc-108080; Santa Cruz Biotechnology, Inc.) were transduced into another line of HK-2 cells in the same manner as that of LOXL2 shRNA particles and were further incubated with media containing either TGF-β (20 ng/ml, R&D Systems) or vehicle (0.1% 4mM HCl/BSA) after 24 hours of serum starvation.

### Western blot analysis for LOXL2, Smad-related molecules and epithelial-mesenchymal transformation-related molecules

Fresh frozen kidney tissues from mice and HK-2 cells were homogenised and western blotting was performed following the same protocol of our previous work[14]. Radioimmunoprecipitation assay buffer (Biosesang, Inc., Seongnam, Korea) with a protease inhibitor cocktail (Roche Diagnostics, Indianapolis, IN,v USA) was prepared. After lysing the cells in buffer, the samples were centrifuged at 13,000 rpm for 30 min at 4°C. The protein concentration was measured through bicinchoninic acid protein assay kit (Thermo Fisher Scientific) according to the manufacturer’s protocol. When the protein samples (50 μg) were separated by 10% sodium dodecyl sulphate-polyacrylamide gel electrophoresis (SDS-PAGE) for 2 h at 100 V, they were transferred onto a polyvinylidene fluoride membrane and blocked with 3% skim milk for 1 h at room temperature. Primary antibodies were incubated with the membrane overnight at 4°C. The anti-mouse-specific primary antibodies purchased from Cell Signaling Technology (Danvers, MA, USA) included anti-Smad2 (cat. no. 5339; 1:1000), anti-physphorylated-Smad2 (p-Smad2, Ser465/467) (cat. no. 3108; 1:500), anti-Smad3 (cat. no. 9523; 1:1000), anti-phosphorylated-Smad3 (p-Smad3, Ser423/425) (cat. no. 9520; 1:1000), anti-Smad2/3 (cat. no. 8685; 1:1000), and anti-Smad4 (cat. no. 38454; 1:1000). The blocking solution used for the anti-phospho-Smad2 and anti-phospho-Smad3 antibodies contained 5% bovine serum albumin (Sigma-Aldrich). The primary antibodies applied to HK-2 cells were anti-α-smooth muscle actin (α-SMA, cat. no. A5228; 1:500; Sigma-Aldrich), anti-vimentin (cat. no. ab92547; 1:5000; Abcam, Cambridge, MA, USA), anti-E-cadherin (cat. no. 610181; 1:500; BD Biosciences, San Jose, CA, USA), anti-zona occludens (ZO)-1 (cat. no. ab2272; 1:500; Sigma-Aldrich), anti-fibronectin (cat. no. sc8422; 1:1000; Santa Cruz Biotechnology, Inc.), and anti-LOXL2 (cat. no. ab96233; 1:500; Abcam). The membrane was washed with Tris-buffered saline which contains 0.1% Tween-20. It was then incubated with horseradish peroxidase-labelled secondary antibodies (cat. no. sc-2020; 1:5,000; Santa Cruz Biotechnology, Inc.; and cat. no. K4003; 1:5,000; Dako; Agilent Technologies, Inc., Santa Clara, CA, USA) for 1 hour at room temperature. Pierce Enhanced Chemiluminescence Western Blotting Substrate (Thermo Fisher Scientific) was used to visualize protein bands. The membrane was stripped with Restore Western Blot Stripping Buffer (Thermo Fisher Scientific) for 15 min at room temperature, and then it was incubated with an anti-β-actin antibody (cat. no. sc-47778; 1:2,000; Santa Cruz Biotechnology, Inc.). The bands were semi-quantified by densitometry using Image J software (version 1.50i; National Institutes of Health, Bethesda, MD, USA).

### Statistical analysis

Quantitative analysis was performed for the western blot and RT-qPCR results. Vehicle-injected mice, folic acid-injected mice treated with AB0023, and folic acid-injected mice treated with control IgG were compared. Data are expressed as the mean ± standard deviation and compared using the Mann-Whitney U test, one-way analysis of variance, Kruskal-Wallis test, and Wilcoxon signed rank test. The analyses were performed using SPSS version 25 software (IBM-SPSS Inc., Armonk, NY, USA). Differences with p<0.05 were considered statistically significant.

## Results

### LOXL2 inhibition prevented the progression of tubulointerstitial fibrosis in the mouse model

The amount of fibrosis measured by trichrome (Fig 2A) and picro-sirius red staining (Fig 2B) decreased in mice treated with AB0023, compared to the control IgG-treated group (Fig 2C and D). Quantitative measurement of fibrosis by total collagen analysis also showed that fibrosis decreased in mice treated with AB0023 (Fig 2E).

**Fig 2.**
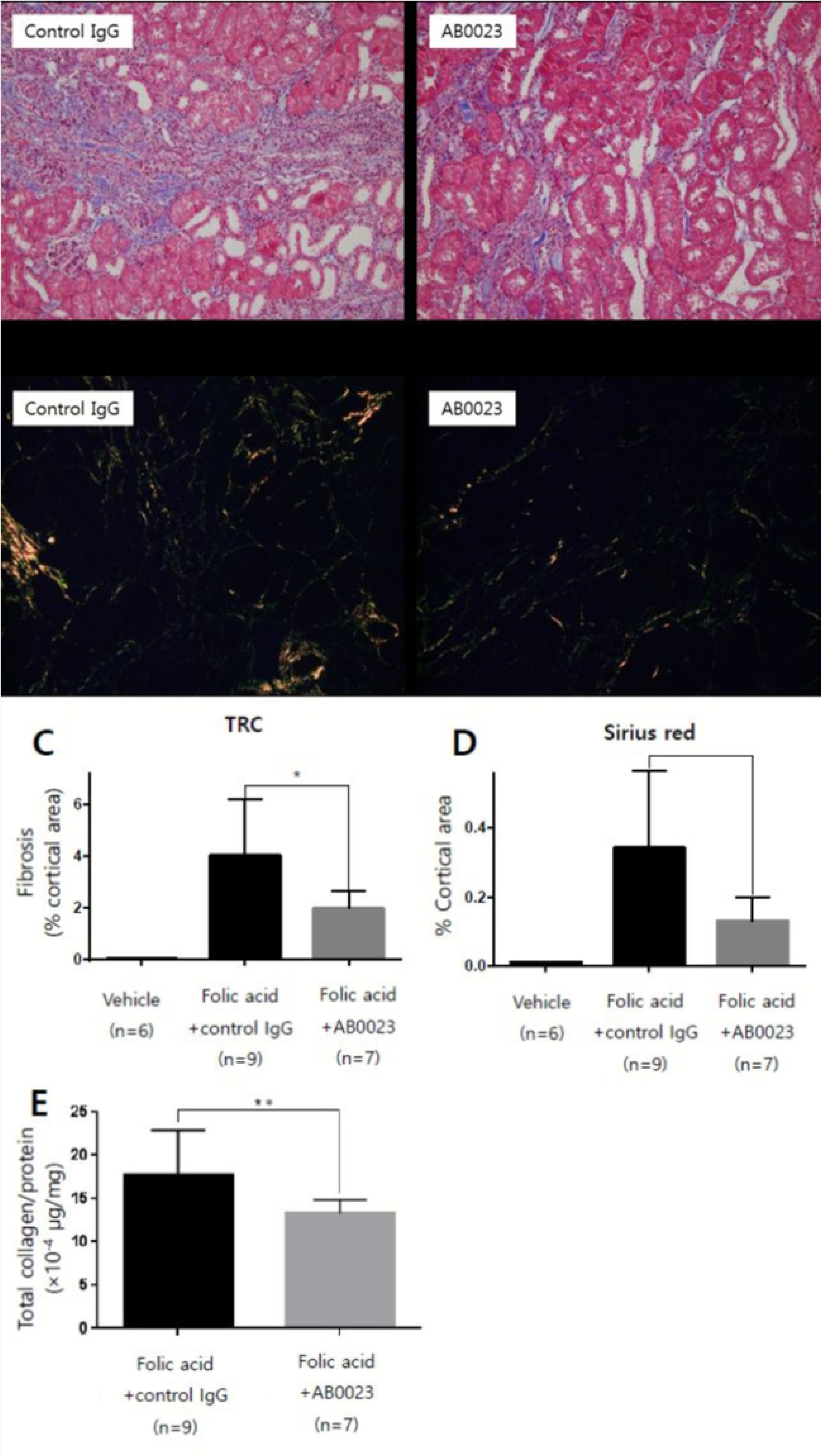
Effects of LOXL2 inhibition on the progression of tubulointerstitial fibrosis. Representative photos of renal cortex of folic acid-injected mice treated with control IgG or AB0023. The amount of fibrosis was measured using Masson trichrome stain (A&C) and picro-sirius red stain under polarized microscopy (B&D). The levels of fibrosis were significantly lower in the AB0023-treated group compared to the control group. The total collagen assay showed similar result (E). *p<0.017, **p<0.05. LOXL2, lysyl oxidase-like 2.

### LOXL2 inhibition may influence the canonical TGF-β/Smad signalling pathway

Smad signaling pathway molecules, including p-Smad3, p-Smad2, and Smad4 exhibited no significant difference with LOXL2 inhibition (Fig. 3). However, the amounts of p-Smad2 and Smad4 tended to decrease in the AB0023-treated group relative to the control group.

**Fig 3.**
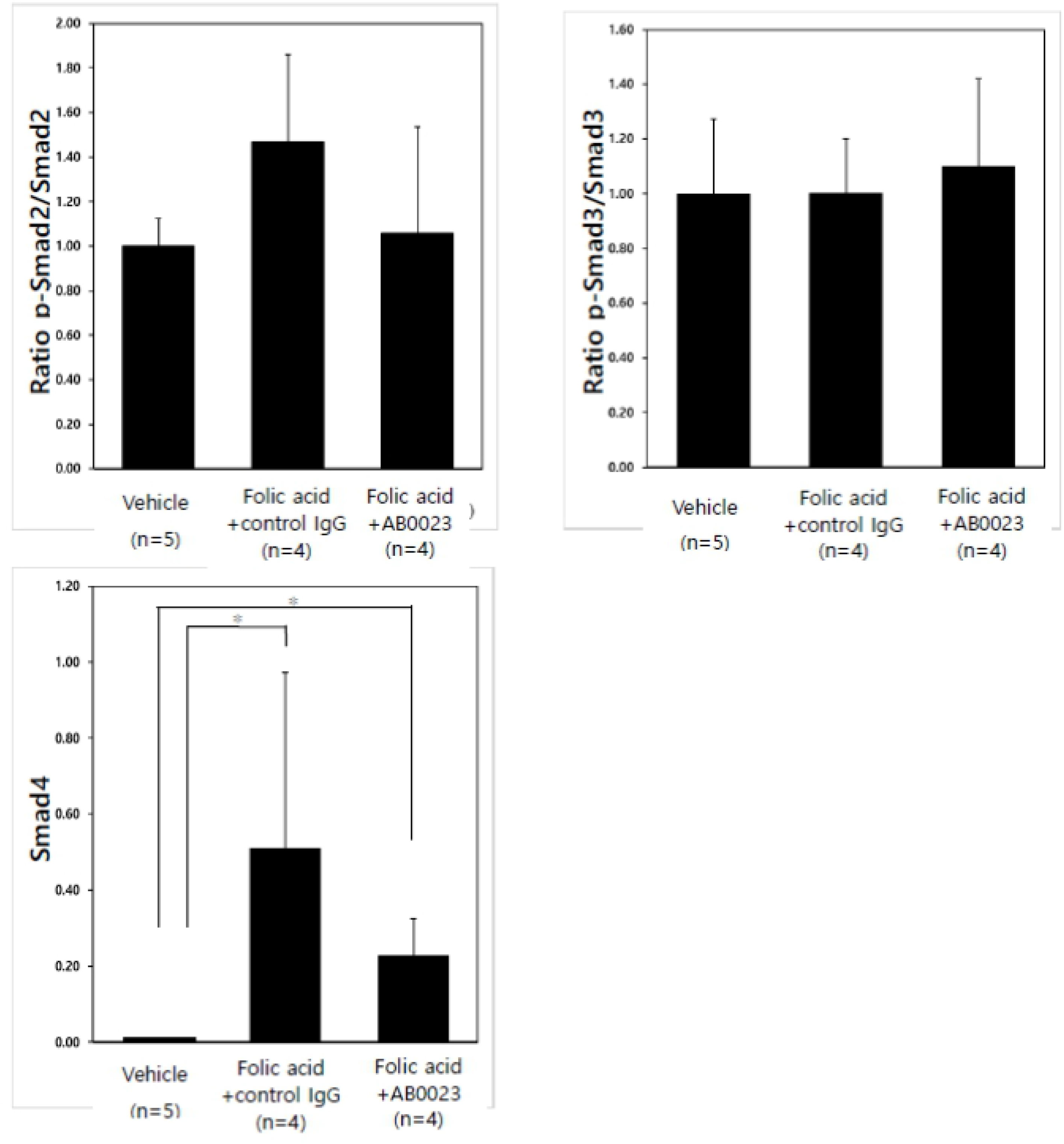
Effects of LOXL2 inhibition on the TGF-β/Smad pathway. Effects of LOXL2 inhibition on the TGF-c/Smad pathway. There was no significant difference in the level of Smad pathway-related molecules, but the amount of p-Smad2 and Smad4 tended to be lower after LOXL2 inhibition. P-values: 0.376, 0.784, and 0.010 for p-Smad2/total Smad2, p-Smad3/total Smad3, and Smad4, respectively.*p<0.017. LOXL2, lysyl oxidase-like 2.

### LOXL2 knockdown in HK-2 cells reduced the expression of some EMT-associated molecules

Transfection of HK-2 cells with LOXL2 shRNA resulted in LOXL2 knockdown (see S1 Fig). In control HK-2 cells transfected with control shRNA, TGF-β treatment (72 hours) reduced the levels of epithelial marker E-cadherin, and increased the levels of mesenchymal markers vimentin and fironectin. Multiple comparison analysis revealed that vimentin level was significantly lower in LOXL2 knockdown cells than in control shRNA-transduced cells after TGF-β treatment (Fig 4). Considering that the level of vimentin increases after TGF-β challenge in control cells, the decreasing trend of the vimentin level in LOXL2 knockdown cells after TGF-β challenge is more meaningful. The epithelial markers ZO-1 and E-cadherin did not show a significant difference after TGF-β treatment in LOXL2 knockdown and control cells, while the level of E-cadherin was markedly decreased by TGF-β challenge.

**Fig 4.**
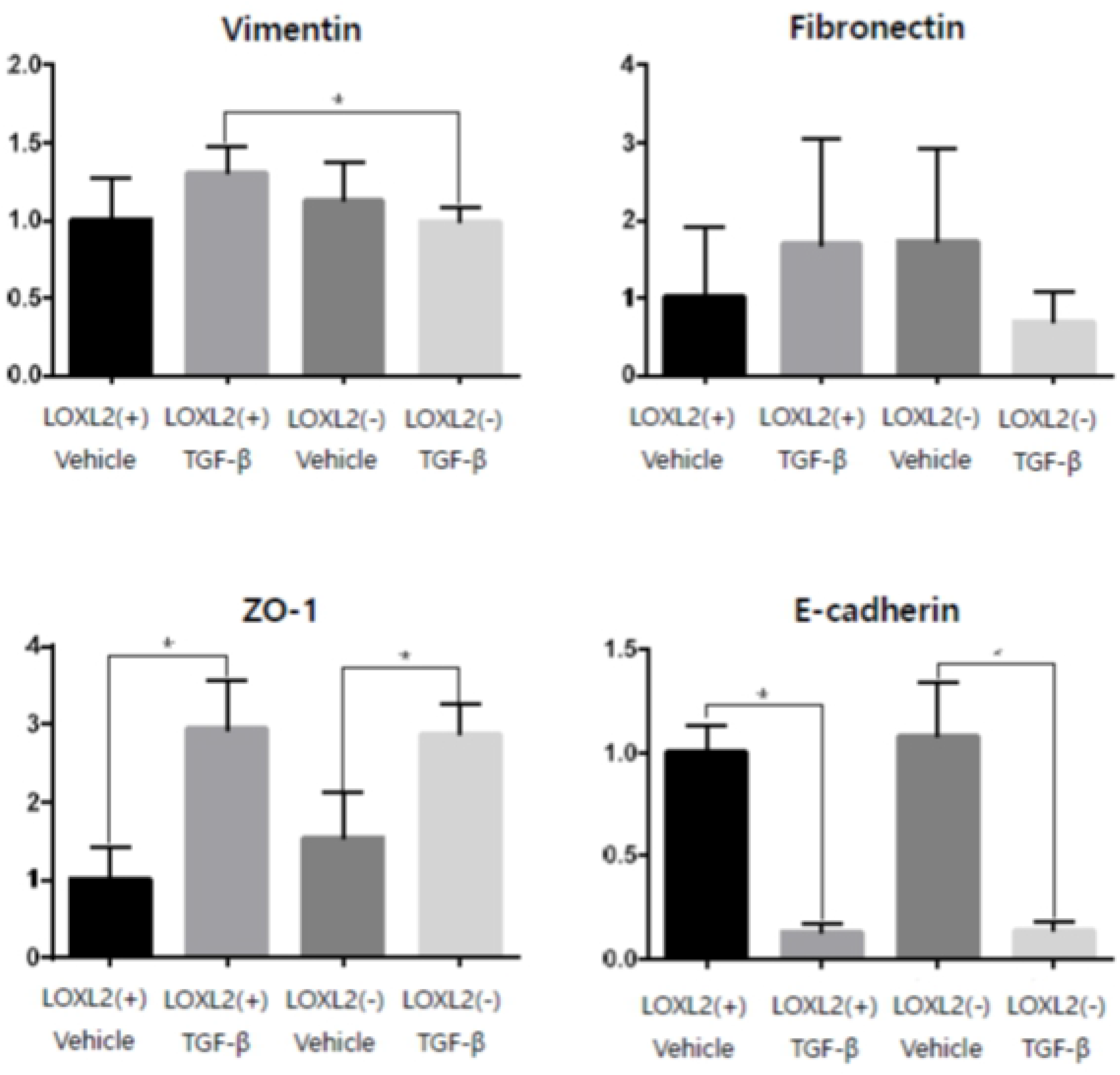
Expression of EMT-associated molecules after TGF-β challenge in LOXL2 knockdown HK-2 cells. HK-2 cells were transfected with either control shRNA virus or LOXL2 shRNA lentivirus to knockdown LOXL2 expression. After 24 hours of serum starvation, both the LOXL2-deficient cells and control cells were incubated with either TGF-β or vehicle for 72 hours. Compared to control HK-2 cells, LOXL2-deficient HK-2 cells showed decreased expression of vimentin after TGF-β challenge. *p<0.05. N=6 for all four groups. LOXL2(+), HK-2 cells transfected with control shRNA; LOXL2(−), LOXL2-deficient HK-2 cells by LOXL2 shRNA transfection; LOXL2, lysyl oxidase-like 2; ZO-1, zona occludens-1.

## Discussion

Inhibition of LOXL2 via AB0023 has been shown to reduce fibrosis in various organs. For instance, AB0023 attenuated postoperative fibrosis in a rabbit model of glaucoma surgery [18]. AB0023 also attenuated tetrachloride-induced hepatic fibrosis in BALB/c mice, and high-dose bleomycin-induced pulmonary fibrosis in C57BL/6 mice [19]. Although a clinical trial of simtuzumab, a humanized form of AB0023, resulted in no significant changes in fibrosis in human immunodeficiency virus- and hepatitis C virus-infected adults, serum samples suggested up-regulation of TGF-β3 and interleukin-10 pathways with treatment, suggesting the future evaluation for clinical trials with simtuzumab after the modulation of TGF-β3 [20]. We previously observed that LOXL2 is expressed in tubular epithelial cells, and presumed a role for LOXL2 in TGF-β-mediated tubulointerstitial fibrosis. This hypothesis is supported by the increased LOXL2 mRNA and protein levels detected in the kidneys of mice with folic acid-induced tubulointerstitial fibrosis [11]. The positive regulatory role of LOXL2 in tubulointerstitial fibrosis is confirmed by the reduction of fibrosis in this folic acid-induced fibrosis mouse model with the inhibition of LOXL2 by the LOXL2-specific antibody AB0023. To the best of our knowledge, this is the first study demonstrating that LOXL2 inhibition leads to attenuated fibrosis in the murine kidney injry model..

In addition to fibrosis, TGF-β is involved in various biological activities, such as cell proliferation, apoptosis, differentiation, autophagy, and the immune response [21]. Thus, it is critical to investigate therapeutic strategies related to the downstream pathways of TGF-β due to the possible adverse effects of directly targeting this cytokine [22].

EMT is a major mechanism that contributes to renal fibrosis in response to multiple molecules, including TGF-β1 [1], connective tissue growth factor [23], and angiotensin II [11], in tubular epithelial cells. However, TGF-β1 is the most potent inducer of EMT [5, 8]. As mentioned above, EMT is a well-described process characterized by a loss of epithelial cell adhesion molecules, such as E-cadherin and ZO-1, *de novo* α-SMA expression and actin filament reorganization, transformation of myofibroblastic morphology, tubular basement membrane disruption, and cell migration/infiltration into the interstitium [5, 8]. However, conflicting results have been reported from *in vivo* studies as tubular cells that have undergone partial EMT relay proinflammatory and profibrogenic signals to the interstitium without directly contributing to the myofibroblast population [24]. Accordingly, the relationship between LOXL2 and the TGF-β pathway *in vivo,* and the TGF-β-mediated relationship between LOXL2 and EMT *in vitro*, were investigated in this study. Thus, canonical pathway-related molecules were studied *in vivo* and markers expressed by tubular cells during EMT were studied *in vitro*. The lack of significant differences in the level of Smad molecules after LOXL2 inhibition in this study may be due to the lapse of time between folic acid injection and analysis. Murine kidneys were harvested at 4 weeks after this injury, by which time fibrogenesis could have already been completed. Stallons et al. reported that TGF-β1 and α-SMA levels increased until 6 days after folic acid injection, and gradually decreased afterwards in a similar experiment where a 250 mg/kg dose of folic acid was injected intraperitoneally [16]. Tang et al. reported that after the administration of high glucose doses, the expression of p-Smad2 and p-Smad3 increased in HK-2 cells for 30 to 60 min and 30 to 120 min, respectively, before decreasing gradually [25]. Our study differed from this experiment; in particular, the time between intervention and injury was substantially longer than that in previous studies. A more rapid analysis of Smad molecules after folic acid injection may have revealed a more pronounced change in their expression in this study.

Experiments on HK-2 cells *in vitro* after TGF-β challenge revealed less increase in the myofibroblast marker vimentin in LOXL2 knockdown cells. These data indicate that LOXL2 plays a regulatory role in EMT. Other studies have shown no reduction in epithelial markers, such as E-cadherin, with LOXL2 inhibition after TGF-β challenge, while a significant reduction of E-cadherin was observed here. However, cell types and experimental methods used in this study differ from those reported previously [12, 13, 26]. Although E-cadhrin and ZO-1 level remained insignificant, it can be inferred that LOXL2 might be related to EMT pathway based on significant change of vimentin. Further studies are warranted to elucidate the mechanisms underlying the LOXL2-EMT-related pathway.

In conclusion, inhibition of LOXL2 ameliorates renal fibrosis. LOXL2 is associated with TGF-β-mediated tubulointerstitial fibrosis and EMT. Improved understanding of the role of LOXL2 in the kidney may illuminate the pathophysiology of tubulointerstitial fibrosis and glomerulosclerosis, and potentially lead to the discovery of novel therapeutic targets for treating these conditions.

## Supporting information

**S1 Fig. Transfection of HK-2 cells with LOXL2 shRNA resulted in LOXL2 knockdown.**

HK-2 cells were transfected with either LOXL2 shRNA lentivirus to knockdown LOXL2 expression, or control shRNA virus. After 24hours of serum starvation, both the LOXL2-deficient cells and control cells were incubated with either TGF-β or vehicle for 72 hours. Whereas LOXL2 increased in control HK-2 cells after TGF-β challenge, LOXL2-deficient cells showed no significant difference in LOXL2 level. *p<0.05. LOXL2(+), HK-2 cells transfected with control shRNA; LOXL2(−), LOXL2-deficient HK-2 cells by LOXL2 shRNA transfection; LOXL2, lysyl oxidase-like 2.

